# Genetic regulatory effects modified by immune activation contribute to autoimmune disease associations

**DOI:** 10.1101/116376

**Authors:** Sarah Kim-Hellmuth, Matthias Bechheim, Benno Pütz, Pejman Mohammadi, Yohann Nédélec, Nicholas Giangreco, Jessica Becker, Vera Kaiser, Nadine Fricker, Esther Beier, Peter Boor, Stephane Castel, Markus M. Nöthen, Luis B. Barreiro, Joseph K. Pickrell, Bertram Müller-Myhsok, Tuuli Lappalainen, Johannes Schumacher, Veit Hornung

## Abstract

The immune system plays a major role in human health and disease, and understanding genetic causes of interindividual variability of immune responses is vital. We isolated monocytes from 134 genotyped individuals, stimulated the cells with three defined microbe-associated molecular patterns (LPS, MDP, and ppp-dsRNA), and profiled the transcriptome at three time points. After mapping expression quantitative trait loci (eQTL), we identified 417 response eQTLs (reQTLs) with differing effect between the conditions. We characterized the dynamics of genetic regulation on early and late immune response, and observed an enrichment of reQTLs in distal *cis*-regulatory elements. Response eQTLs are also enriched for recent positive selection with an evolutionary trend towards enhanced immune response. Finally, we uncover novel reQTL effects in multiple GWAS loci, and show a stronger enrichment of response than constant eQTLs in GWAS signals of several autoimmune diseases. This demonstrates the importance of infectious stimuli modifying genetic predisposition to disease.

## Main Text

An increasingly popular approach to identify genetic factors affecting interindividual variation in immune response is mapping expression quantitative trait loci (eQTLs) – variants that associate to gene expression – and to identify so-called response eQTLs (reQTLs) where the eQTL effect differs between immune stimuli^1-6^. Such genetic variants can impact the transcriptional response to infection, and also represent genetic effects that are modified by the infectious environment via gene-by-environment interactions. In this study, we create a data set of a large number of immune stimulus conditions, with monocytes activated with microbial ligands for three different pattern recognition receptor (PRR) families at two different time points, next to the baseline condition.

To examine the time course of innate immune responses, we first profiled gene expression in monocytes of five individuals using Human HT-12 v4 Expression BeadChips (Illumina) at six time points after stimulation with three prototypical microbial ligands: Lipopolysaccharide (LPS) was used to activate TLR4, muramyl-dipeptide (MDP) to stimulate NOD2, and 5’-triphosphate RNA (RNA) to activate RIG-I. Hierarchical clustering revealed early differentially expressed (DE) genes at 45 and 90 minutes after stimulation and late DE genes between 3 and 24 hours (**Supplementary Fig. 1**). For the full eQTL cohort, we analyzed primary monocytes isolated from 134 healthy male individuals (185 before quality control), which were either untreated (baseline) or stimulated with the same three pathogen-derived stimuli, and gene expression was profiled after 90 minutes and 6 hours. All donors were SNP genotyped using Illumina HumanOmniExpress BeadChips (**Fig.1a**). In a previous study^1^ we have analyzed a subset of the data consisting of baseline and 90 minutes LPS-stimulated monocytes in this cohort.

**Figure 1.**
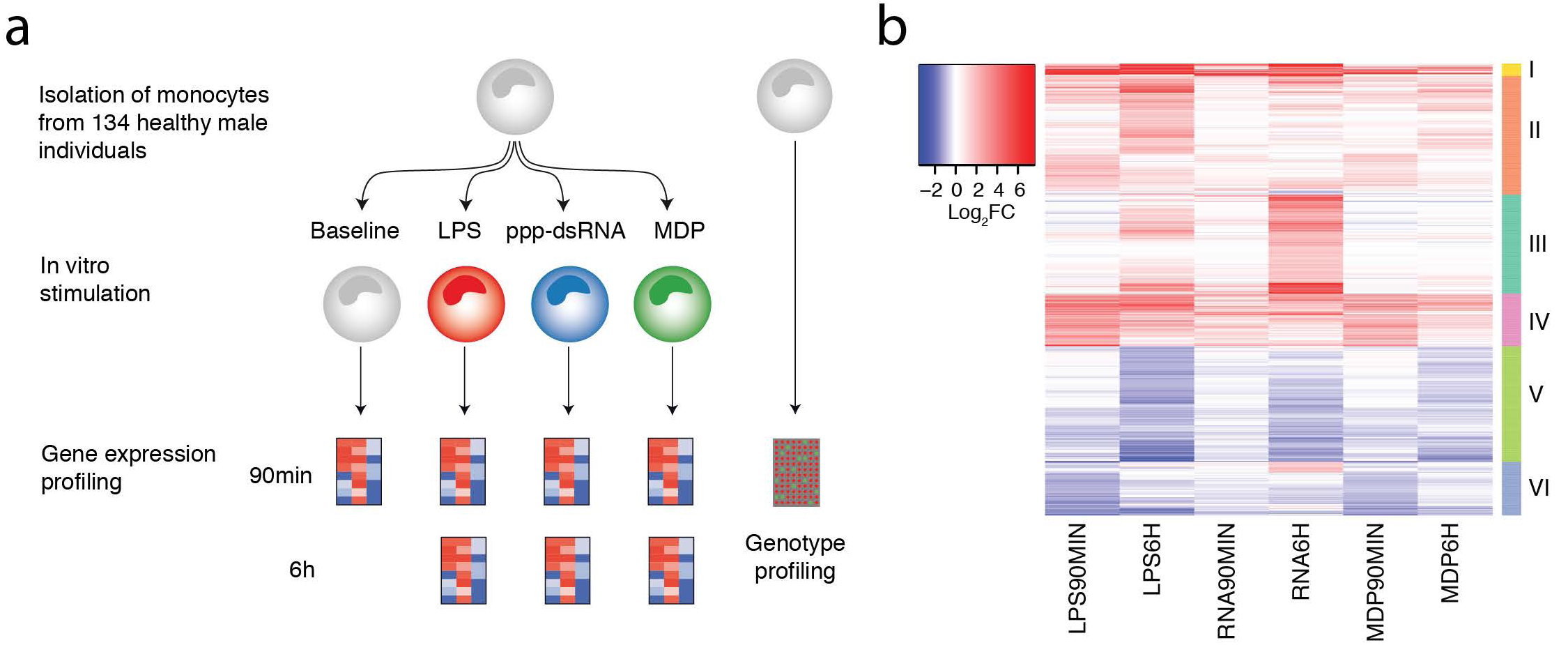
Overview of the study. **(a)** Step-wise experimental design to identify genetic effects on immune response in human monocytes. 1) Isolation and stimulation of primary monocytes from 134 individuals, 2) Transcriptome measurement of the entire cohort at two time points (90min and 6h) after stimulation, 3) Genotype profiling to map immune response eQTLs. **(b)** Mean mRNA profiles of differentially expressed genes (log_2_-fold change > 1, FDR 0.001) of 134 individuals between baseline and each of the six stimulated conditions. Genes are hierarchically clustered into six distinct expression patterns (see **Supplementary Table 1** for a full list of the differential expression and enriched pathways of each cluster).

First, we studied the gene expression response to immune stimulation. Principal component analysis of the gene expression data identified seven distinct groups corresponding to each treatment and time point (**Supplementary Fig. 2**). Differential expression analysis of genes expressed in at least one of the seven conditions showed the highest number of DE genes under late LPS response, and lowest under early RNA stimulation (**Supplementary Fig. 3, Supplementary Table 1**). These genes form six clusters with similar response patterns across time points and conditions (**Fig. 1b, Supplementary Table 1**), and with gene ontology (GO) enrichments corresponding to relevant immunological pathways (**Supplementary Table 1**). Furthermore, immune responsive genes showed a significantly greater and a more diverse distribution of interindividual variance than all expressed genes already in the baseline condition, with a further increase upon stimulation (**Supplementary Fig. 4**). These analyses of gene expression patterns in a population scale provide a highly robust and comprehensive data set of innate immune responses and their interindividual variation upon diverse microbial ligands and multiple time points.

In order to study genetic variation affecting gene expression levels, we performed eQTL mapping for all seven conditions, defining *cis* eQTLs within 1 Mb interval on either side of an expression probe at a false discovery rate (FDR) of 5%. We identified 717 to 1,653 genes with an eQTL in each condition (**Fig. 2a, Supplementary Table 2**). The eQTLs from conditions analyzed in previous studies^3,4^ had a high degree of replication, demonstrating the robustness of our data set (**Supplementary Fig. 5a; Methods**). We provide a user-friendly access to our results via the ImmunPop QTL browser (http://immunpop.com/kim/eQTL).

**Figure 2.**
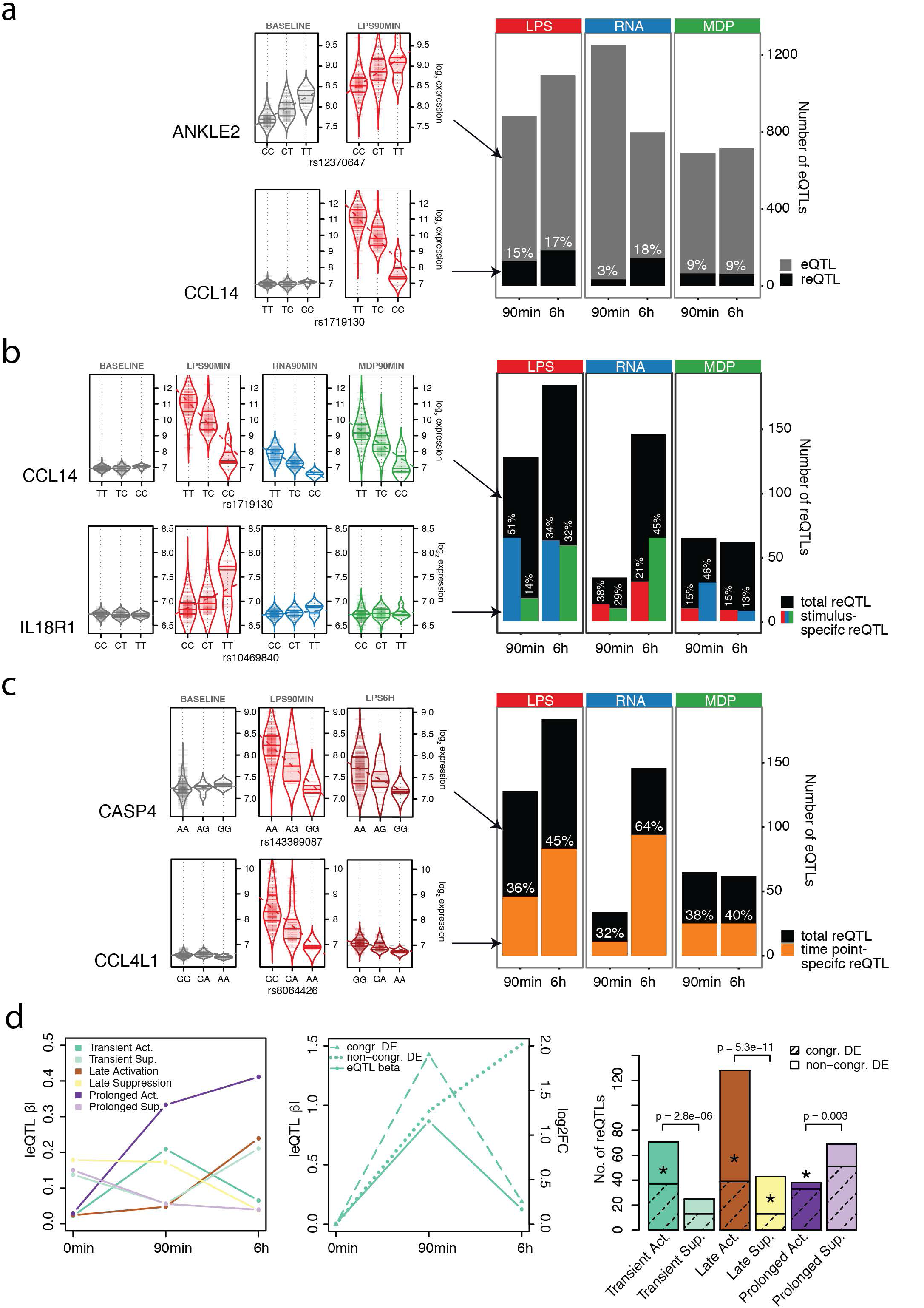
Immune response eQTL study in human monocytes. **(a)** Total numbers of *cis* eQTLs and proportions of reQTLs of LPS-treated (LPS), ppp-dsRNA-treated (RNA) and MDP-treated (MDP) monocytes at 90min and 6h after stimulation. eQTLs include all genes with a significant genetic association in each stimulated condition, and reQTLs are a subset that show a significant difference of the regression slope between untreated and stimulated monocytes, with violin plots shown as examples. The untreated condition has 1,653 eQTLs that are not shown in the barplot. **(b)** Numbers of reQTLs and proportions of treatment-specific reQTLs where the regression slope of the tested treatment is different from the slope of the other two treatments within the same time point, with violin plots shown as examples and the color of bar indicating the treatment that was tested. **(c)** Numbers of reQTLs and proportions of time point-specific reQTLs where the regression slope of the tested time point is different from the slope of the other time point within the same treatment, with violin plots shown as examples. **(d)** reQTLs were divided into six subsets according to their temporal activity (see Methods). Average of absolute eQTL effect sizes per category is shown on the left panel. The middle panel illustrates a reQTL example with congruent differential expression (DE) (dashed line) or non-congruent DE (dotted line) of the eGene. reQTL distribution to different categories is shown in the right panel, where the shaded portion illustrates the proportion of reQTLs with congruent DE of the eGene and asterisks represent the significance of enrichment of reQTLs with congruent DE of the eGene (Fisher’s exact test *p < 0.05). The p-values above the bars indicate the significance between of active and suppressive types (binomial test).

To identify eQTLs that differ between stimuli, we used a beta-comparison approach, comparing the regression slopes of an eQTL under baseline (*β*_baseline_) vs. stimulated (e.g. *β*_LPS90min_) in a z-test, with reQTLs defined as having Bonferroni corrected *p* < 0.05 (see Methods). This approach is highly consistent with a previously used method where differential expression is used as the quantitative trait (**Supplementary Fig. 5b**), but provides more flexibility for comparing several conditions. This analysis revealed that 3-18% of our *cis* eQTLs in each condition are reQTLs (**Fig. 2a, Supplementary Table 2**). Genes with a reQTL showed GO enrichment in immune pathways (**Supplementary Fig. 6a**), and include key genes of protein-protein interaction networks such as MAP kinases, IRF transcription factors, chemokines, and chemokine receptors (**Supplementary Fig. 6b, c, d**), demonstrating the relevance of genetic interindividual variation in the innate immune system.

Next, to analyze treatment and time point specificity of reQTLs we performed pairwise comparisons of regression slopes across treatments and time points, respectively. This revealed that 13-51% of reQTLs were treatment-specific when compared to the other two stimuli of the same time point with marked differences depending on which stimulus-pair was tested (**Fig. 2b**). We also observed a large proportion of time point-specific reQTLs (32-64%) suggesting a highly dynamic genetic regulation in immune response (**Fig. 2c**). Of note, the number of identified reQTLs per condition, as well as time point- and stimulus-specific reQTLs, were correlated to the number of differentially expressed genes (**Supplementary Fig. 7a, b**). Thus, differential expression analysis in a small number of samples can be used to select the conditions that maximize novel reQTL discovery in a population-scale study.

To obtain better insights into the dynamic link between reQTLs and differential expression upon immune stimulation, we classified reQTLs to those with early transient, late, and prolonged effects (see Methods). We find that active reQTLs that are absent under baseline and active under stimulus are more common and have higher effect sizes than suppressive reQTLs where a baseline eQTL is lost under stimulus (**Fig. 2d, Supplementary Fig. 7c**). Interestingly, active reQTLs are typically more dynamic with early transient or late effects, whereas suppressive reQTLs are more often prolonged, extending over both time points. Next, we analyzed whether the temporal dynamics of reQTLs correspond to dynamics of differential expression. A highly congruent pattern would indicate a major role of genetic interindividual variation in reQTL genes across the gene’s temporal response to stimulus, whereas a divergent pattern could suggest recruitment of additional expression response mechanisms independent of the regulatory effect of the reQTL variant. The proportion of reQTL genes with congruent differential expression ranged between 30-87% for different classes of dynamic reQTLs (**Fig. 2d, Supplementary Fig. 7d, Methods**) with significant enrichment of congruent pattern in 4 out of 6 groups (*p* < 0.05 in Fisher’s exact test of each group vs all others). This indicates that reQTLs are relevant regulators of differential expression but additional regulatory mechanisms are involved in shaping the transcriptional response of reQTL genes. Altogether, our analysis of temporal reQTLs sheds light on mechanisms of the highly dynamic immune response, and the role of genetic variants in it.

To further characterize the genetic variants underlying the total of 417 reQTLs across all treatment conditions, we defined a set of 677 constant eQTLs (ceQTL) that display no change in regression slope across all conditions (nominal *p* > 0.05) (**Fig. 3a, Supplementary Fig. 8a**). Functional annotation enrichment and fine mapping analyses by fgwas^7^ revealed that reQTLs were more enriched in promoter flanking regions, CTCF binding sites and enhancer regions, while constant eQTLs were more common in promoter regions, 3’ and 5’UTRs, and regions downstream of TSS (**Fig. 3b, Supplementary Fig. 8b**). While reQTL enrichment has been previously described for some transcription factors^5,6^, and annotations of condition-specific epigenomic marks and tissue-specific eQTLs have been described^8,9^, our results are to our knowledge the first demonstration of environmentally responsive eQTLs being enriched in distal *cis*-regulatory elements.

**Figure 3.**
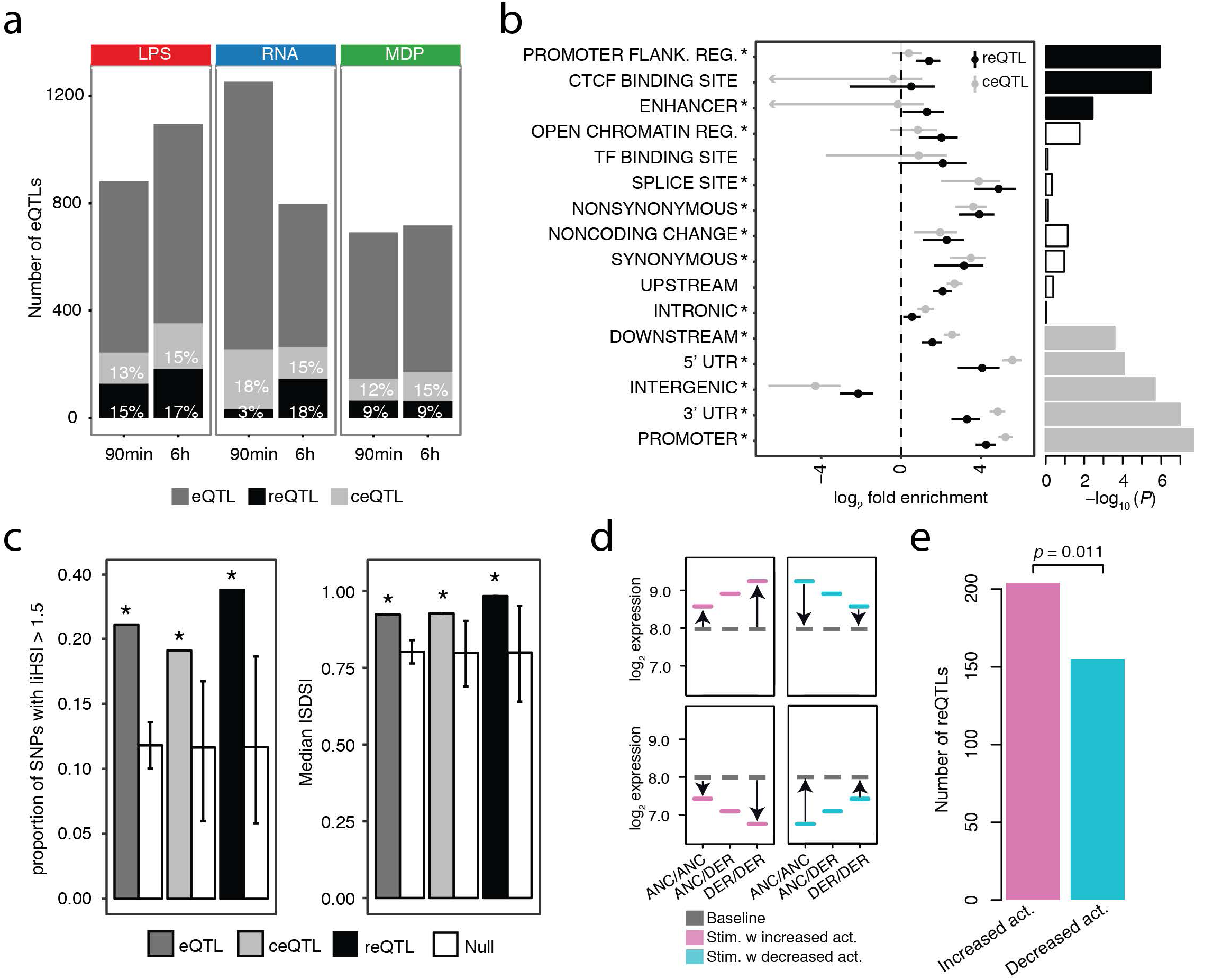
Functional annotations and signs of natural selection in reQTLs. **(a)** Total numbers of *cis* eQTLs, and proportions of reQTLs and constant eQTLs (ceQTL) that have similar regression slopes across all conditions. Examples of a ceQTL and reQTL are shown in Supplementary Fig. 8a. (**b**) Forest plot of enrichment estimates of reQTL and ceQTL signals for each functional annotation with 95% confidence intervals (see also Supplementary Fig. 5b). Bar plot shows the enrichment of the single most likely causal SNP per locus after fine mapping. The solid bars indicate significant enrichments after Bonferroni correction. **(c)** Signal of positive selection measured as the proportion of variants with high |iHS| (left panel), and median |SDS| (right panel), using the variant with the maximum value from each locus across all SNPs in high LD (*r^2^* > 0.8). Genome-wide null sets of variants matched to eQTL, ceQTL or reQTL were generated by resampling 10,000 sets of random SNPs that matched for MAF and LD (white bars). Error bars indicate minimum and maximum of the null distribution, and asterisks indicate the significant enrichment compared to the null (permutation test *p* < 10^−4^). **(d)** Illustration of reQTLs where the derived allele causes an increase (left panel) or decrease (right panel) in response amplitude compared to the ancestral allele. The increase or decrease of the response amplitude can be in both directions, e.g. reQTLs that amplify the induction or amplify the suppression of a gene are both considered as reQTLs with “increasing activity” of the derived allele and reQTLs that weaken the induction or suppression of a gene are both considered as reQTLs with “decreasing activity” of the derived allele. **(e)** Numbers of reQTLs with increased or decreased activity across all stimulated conditions, with a p-value from a binomial test.

Given that the innate immune system is the first line of defense in early interaction between the host and the microbe, we asked if selective pressures that are exerted by microorganisms on the host genome can be detected in reQTLs. Consistent with previous reports^5,6^, we detected a signal of increased positive selection in eQTLs, ceQTLs, and reQTLs using the integrated haplotype score^10^ (iHS; permutation test *p* < 10^−4^, **Fig. 3c**, left panel) and the singleton density score^11^ (SDS; permutation test *p* < 10^−4^, **Fig.3c**, right panel), comparing each eQTL class to a genome-wide null set of variants matched for minor allele frequency (MAF) and linkage disequilibrium (LD). Next, we examined the direction of the effect of the derived allele, dividing reQTLs into two groups (**Fig. 3d**): 1) reQTLs where the derived allele causes an increase in response amplitude compared to the ancestral allele (e.g. ancestrally upregulated genes are further upregulated among derived allele carriers), and 2) reQTLs where the derived allele causes weakening or even silencing of immune response compared to the ancestral allele. Interestingly, across all treatments the reQTLs with stronger expression response by the derived allele were more common (binomial *p* = 0.011 across all conditions; **Fig. 3e, Supplementary Fig. 9**). This suggests an evolutionary trend towards enhanced immune response, which might reflect an arms race of the host immune system and invading pathogens.

Given the central role of inflammation in many diseases, we examined reQTLs as a potential mechanism underlying genetic associations to complex diseases, discovered by genome-wide association studies (GWAS). First, we identified individual GWAS loci that are likely to share a causal variant with an reQTL in the same locus. We used the coloc^12^ method on summary statistics of 33 GWAS traits (**Supplementary Table 3**) and our reQTL data. This analysis provided four loci with strong evidence (PP3 + PP4 ≥ 0.90 and PP4/PP3 ≥ 3) of reQTLs sharing the same causal variant with a GWAS trait (**Fig. 4a,b** and **Supplementary Table 3**). In the chromosome 9 locus associated with HDL^13^ and total cholesterol levels^13^, the eQTL effect for *TTC39B* can be detected at baseline levels, but the increasing effect size upon immune stimulation indicates a possible novel immunological component of TTC39B’s role in the etiology of atherosclerosis. In the *IL18R1* locus associated to celiac disease^14^ (**Fig. 4a**) and the *KLF6* locus associated to schizophrenia^15^ (**Fig. 4b**), the eQTL effects are only present under immune stimulation and would not be discovered in baseline monocytes. Conversely, in the *RNMD1* locus associated to age at menarche^16^, the baseline eQTL effect is diminished upon immune activation. As summary statistics are only available for the minority of GWAS traits, we also identified 29 reQTL genes where the top variant is in high LD (*r^2^* > 0.8) with a disease-associated SNP listed in the GWAS catalog^17^ (**Fig. 4c, Supplementary Table 4**), which may indicate shared causal variants albeit with less certainty than coloc analysis. For ten of these reQTL genes the eQTL was absent under baseline condition (*P*_baseline_ > 0.01), including novel reQTL genes such as *APOL2* potentially associated with glomerulosclerosis, *PTGER4* with allergy, and *PIP4K2A* with acute lymphoblastic leukemia. These results do not exclude other possible mechanisms in other cell types or conditions, but the reQTL analysis discovers novel potential causal genes for individual GWAS loci with an effect that is potentially modified by infections.

**Figure 4.**
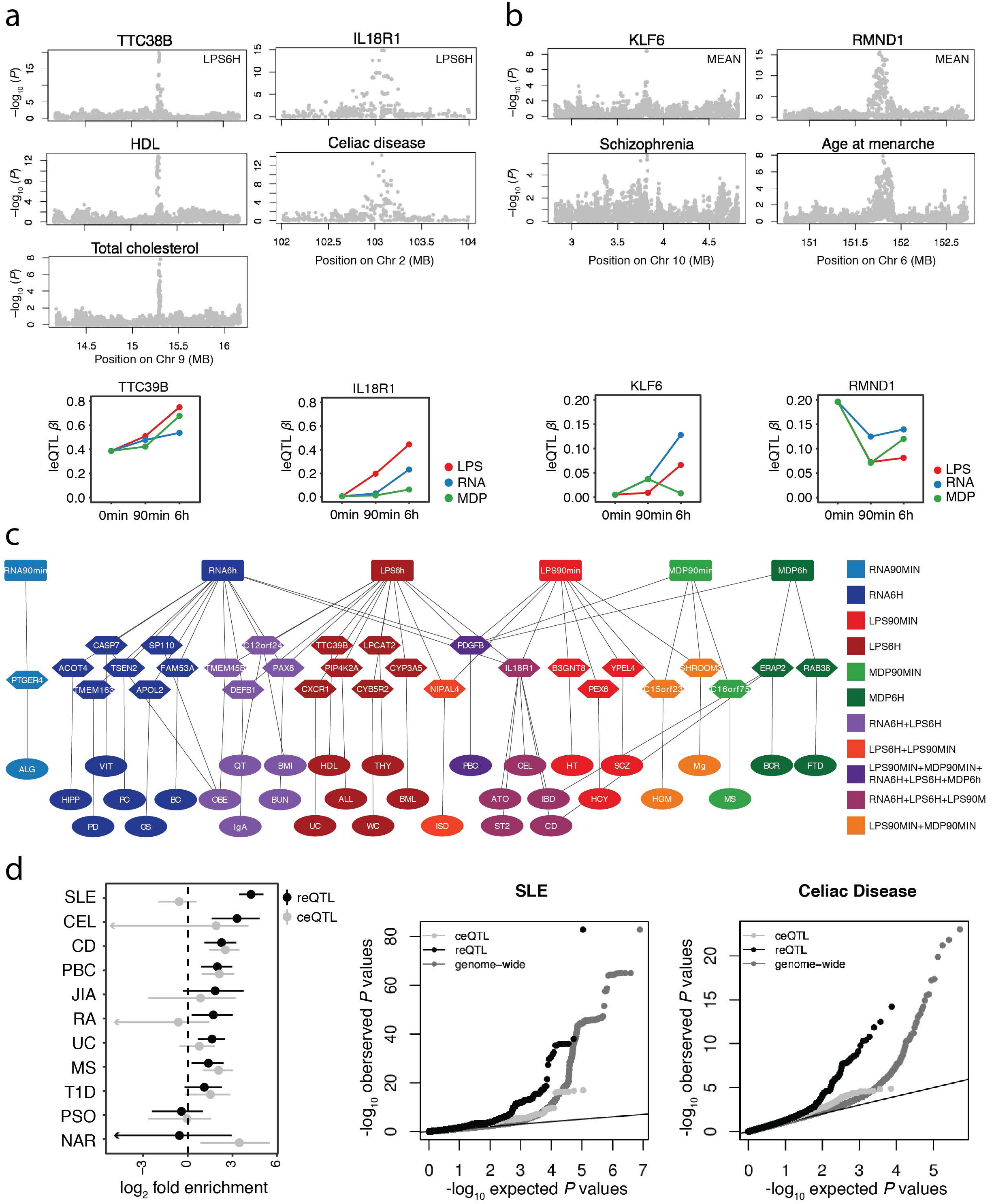
The role of reQTLs in GWAS. **(a)** Manhattan plots of eQTL (top panels) and disease (middle panels) p-values in colocalized loci. The bottom panels show the dynamics of corresponding eQTL effect sizes in different conditions. **(b)** Two additional GWAS loci colocalize when the mean of gene expression across all seven conditions is used to map eQTLs (see Methods). **(c)** Overlap of GWAS SNPs that are in high LD (*r^2^* >0.8) with reQTLs in monocytes with disease phenotypes connected to reQTL genes and corresponding immune stimulations. See Supplementary Table 4 for trait abbreviations. **(d)** Genome-wide enrichment of reQTL and ceQTL associations in autoimmune GWAS with 95% confidence intervals (left panel), and Quantile-quantile (Q-Q) plots for SLE (middle panel) and Celiac disease (right panel). See Supplementary Fig. 10b for additional Q-Q plots, and Supplementary Fig. 11 and Supplementary Fig. 12 for results of non-autoimmune traits.

Finally, to quantify the role of reQTLs in the genome-wide genetic architecture of different complex traits, we analyzed the enrichment of reQTLs and ceQTLs in GWAS signals of eleven autoimmune traits (**Supplementary Table 3**) using fgwas (**Fig. 4d, Supplementary Fig. 10a**), confirmed by Q-Q plots (**Fig. 4d, Supplementary Fig. 10b**) analogously to Li et al.^18^. Interestingly, in seven out of eleven traits reQTLs had a significant enrichment, whereas ceQTLs were enriched in only three of these seven traits, and narcolepsy (NAR) was the only trait significantly enriched for ceQTLs but not for reQTLs. Most notably, systemic lupus erythematosus (SLE) GWAS signals^19^ were very strongly and significantly enriched among reQTLs with no enrichment in ceQTLs, suggesting that innate immune response to pathogens may be a particularly important environmental modifier of genetic predisposition to SLE, while playing a smaller role in the genetic architecture of e.g. psoriasis and type 1 diabetes. While some non-autoimmune traits showed an eQTL enrichment, none were significantly enriched among reQTLs but not among ceQTLs (**Supplementary Fig. 11, Supplementary Fig. 12**). These results indicate a substantial, disease-specific role of environmental interactions with microbial ligands in genetic risk to complex autoimmune diseases. While tissue-specificity of molecular effects of GWAS variants is increasingly appreciated and analyzed^9^, our results suggest that innate immune stimulation is a key cellular state to consider in future eQTL studies as well as in targeted functional follow-up of GWAS loci.

In this study we analyzed interindividual variability of immune response in activated monocytes and characterized genetic variants that influence the response to pathogen components. Unlike previous studies, we analyze various ligands under multiple time points, and provide a more comprehensive picture of the role of genetic variation in innate immunity. Our analysis sheds light on the dynamics of immune response and reQTLs, the genomic elements underlying *cis* eQTLs responding to environmental stimuli, the evolution of immune response, and the key role of immune activation as a modifier of genetic effects especially in autoimmune diseases.

Several important aspects of genetic regulatory variants affecting transcriptional immune response remain to be addressed by other studies. RNA-sequencing allows increased power and identification of splicing QTLs^5,18,20^, and additional epigenomic assays can provide insight into genomic mechanisms of transcriptome response^21^. Increasing sample sizes would provide better power and allow exploration into rare *cis*-eQTL variants^22,23^ and comprehensive *trans* eQTL mapping. Finally, while our study includes more immune stimuli and time points than previous analyses, it is essential to further expand the number of conditions and cell types involved in innate and adaptive immunity in reQTL studies, and advance their joint analysis. The ImmunPop QTL browser that includes our data provides a step towards this direction.

Taken together, our comprehensive characterization of reQTLs provide novel insights into the genetic contribution to interindividual variability and its consequences on immune-mediated diseases. These results support a model where genetic risk for disease can sometimes be driven not by static and uniform malfunction but rather by failure to respond properly to an environmental stimulus. This emphasizes the importance of context-specific genetic regulation in human traits.

## Methods

### Sample collection, isolation and stimulation of CD14+ monocytes

In total, 185 healthy male volunteers of German descent were recruited. The study was approved by the institutional review board of the University of Bonn. All volunteers were between age 18 and 35 (mean 24). Peripheral blood mononuclear cells (PBMC) were obtained by Ficoll-Hypaque density gradient centrifugation of heparinized blood. Monocytes were isolated by MACS using CD14-microbeads (Miltenyi Biotec) as previously described^1^. The purity of isolated monocytes was ≥ 95%. RPMI 1640 (Biochrom) supplemented with 10% heat-inactivated FCS (Invitrogen), 1.5 mM L-glutamine, 100 U/ml penicillin, 100 μg/ml streptomycin (all Sigma-Aldrich) and 10 ng/ml GM-CSF (ImmunoTools) was used to culture cells in 96-well round bottom wells at a density of 250,000 cells/well in 100 μl overnight. Cell viability after overnight incubation was > 85%. Cells of each volunteer were divided to subsets that were either untreated or treated with 200 ng/ml ultrapure LPS from Escherichia coli (Invivogen), 100 ng/ml L18-MDP (Invivogen) or 200 ng *in vitro* transcribed 5’-ppp-dsRNA transfected with Lipofectamine 2000. Based on the pilot study described in Supplementary Fig. 1 and in the Supplementary Information, cells were lysed in RLT reagent (Qiagen) after 90 min or 6 hours and stored at −80°C. C-reactive protein (CRP) levels were measured to exclude samples with elevated CRP levels. After applying stringent quality control and clinical exclusion criteria (Nonsmoker, no infection or vaccination 4 weeks prior to blood withdrawal, CRP < 2.5 mg/dl, monocyte purity ≥ 95%, monocyte survival > 85%), samples from 134 individuals were further processed.

### RNA extraction

RNA was extracted from lysed cells using the AllPrep 96 DNA/RNA Kit from Qiagen. RNA concentrations were determined using NanoDrop (PeqLab) and a subset of samples was additionally checked for degradation in a Bioanalyzer (Agilent Technologies).

### Gene expression analysis

The Illumina TotalPrep-96 RNA Amplification Kit (Life Technologies) was used for amplification and biotinylation of RNA. Subsequent array-based gene expression analysis was performed on Illumina’s Human HT-12 v4 Expression BeadChips (Illumina) comprising 47,231 probes. Expression profiles were quantile normalized, and only probes, which showed a *P*_detection_ < 0.01 in at least ten samples across all conditions were analyzed. Batch effects were removed using the R packages ComBat^24^ and sva^25^. Probes with an interindividual standard deviation > 5 were set to NA. Probes that matched to multiple positions in the human genome and probes mapping to non-autosomal chromosomes were excluded from further analysis. Finally, probes were removed if SNPs within a probe showed an eQTL effect to the respective gene, resulting in 18,988 probes (13,207 genes) for statistical analyses.

To determine the number of differentially expressed genes, the probe with the best *P*_detection_ across all conditions was used and differential expression (log2-fold change > 1, FDR 0.001) was computed using the linear modeling-based approach implemented in the Bioconductor limma package^26^. Genes differentially expressed in at least one condition were grouped into six distinct clusters corresponding to genes with similar response pattern using hierarchical clustering. Over-representation of Gene Ontology terms in these clusters of differentially expressed genes were assessed using hypergeometric-based tests implemented in the R package GOstats^27^. Genes that were expressed in our monocyte data were used as background set in all enrichment analyses. Only enrichments significant at FDR of 0.05 are reported in Supplementary Table 1.

### DNA extraction

Genomic DNA was extracted from 10 ml blood using Chemagic Magnetic Separation Module I (PerkinElmer Chemagen) according to the manufacturer’s instructions. DNA was quantified by NanoDrop (PeqLab).

### DNA genotyping and imputation

Genotyping was conducted on the Illumina’s HumanOmniExpress BeadChips comprising 730,525 SNPs. After quality control (*P*_HWE_ > 10^−5^, call rate > 98%, MAF > 5%), a total of 579,090 SNPs were available for analysis. Samples showing potential admixture within the multi-dimensional scaling (MDS) analysis were removed. All samples showed a call rate > 99%.

Genotypes were phased with SHAPEIT2^28^ and imputed with IMPUTE2^29^ in 5Mb chunks against the 1000 genomes phase 1 v3 reference panel^30^. Sites with an information score of less than 0.8 or significant departure from Hardy-Weinberg equilibrium (*p* < 10^−5^) or MAF < 5% were excluded from further analysis. Genotype probabilities for all remaining sites were converted into dosage estimates.

### eQTL analysis

As quantitative phenotypes we used absolute expression values of untreated (baseline), LPS-treated (LPS), ppp-dsRNA-treated (RNA) and MDP-treated (MDP) cells. Complete expression profiles of each of the seven conditions (baseline, LPS90min, LPS6h, RNA90min, RNA6h, MDP90min, MDP6h) were available for 134 donors. eQTL mapping was performed for SNPs located within 1Mb of the gene expression probe using FastQTL^31^. Significance of the most highly associated variant per gene was estimated by adaptive permutation with the setting "--permute 100 10000”. Permutation p-values obtained via beta approximation were used to access genome wide significance via Benjamini-Hochberg (FDR < 0.05). Downstream analyses were carried out in R. Network analysis of reQTL genes was performed using the STRING 10.0 database^32^ selecting only interactions that were either experimentally validated or originated from curated databases.

### Replication of eQTLs

We compared our results with two previous reQTL studies. For quantifying eQTL replication with a genome-wide study of monocyte eQTLs^4^, we used Storey’s qvalue R package^33^. The *π_1_* statistic considers the full distribution of association p-values (from 0 to 1) and computes their estimated *π_0_*, the proportion of eQTLs that are truly null based on their distribution. Replication is reported as the quantity *π_1_* = 1 - *π_0_* that estimates the lower bound of the proportion of truly alternative eQTLs.

Lee et al.^3^ used a targeted approach (415-gene signature) to identify eQTLs after LPS, Flu or IFΝβ treatment in dendritic cells. *π_0_* could not be calculated using Lee et al. because less than 10% of eQTL genes in our data were represented in the 415 targeted genes, and thus replication was assessed by the proportion of our eQTLs with nominal significance (*p* < 0.05) in Lee et al.

### Detecting reQTLs by eQTL *β*-comparison

In each condition, we first determined the best eQTL per gene (lead eSNP). Regression coefficient (*β*) and its variance (*σ^2^*) of these eQTLs were calculated for all seven conditions using the linear model function `summary (lm())` in R. We then tested if the regression coefficient of an eQTL was significantly different between two conditions in a z-test:

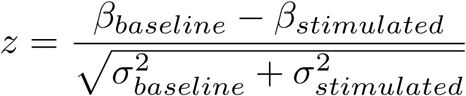

Resulting p-values were corrected for multiple testing using Bonferroni correction (*P*_beta_ < 0.05). Previous reQTL studies^1,3,5^ have used differential expression as a quantitative trait to identify reQTLs (*p*_diff_). We calculated *p*_diff_ for all reQTLs identified by *β*-comparison and used Spearman correlation as a measure of similarity.

To detect treatment specificity of reQTLs, we tested all significant reQTLs of one treatment (e.g. LPS90min) vs the other two treatments of the same time point (e.g. RNA90min and MDP90min) in two separate z-tests. A reQTL was treatment specific if the Bonferroni-corrected p-value in the z-test was < 0.05. To detect time point specificity of reQTLs, for each treatment we tested all significant reQTLs of one time point (e.g. LPS90min) vs the other time point (e.g. LPS6h) in a z-test. Time point specific reQTLs were determined using Bonferroni-corrected p-values (*p* < 0.05). To compare reQTLs with eQTLs that are constitutively active (ceQTL), we defined ceQTLs as eQTLs with *p*_beta_ > 0.05 when testing each of the six stimulated conditions with the baseline condition.

### Characterizing dynamics of reQTLs

To study the dynamics of reQTLs, we encoded as a binary call whether reQTLs had a significant eQTL p-value at each of the three time points or not (e.g. “0- 1-0” codes for “not significant eQTL at 0 min – significant at 90 min – not significant at 6 hours”). If a reQTL was shared between treatments, the treatment with the best p-value was used. This resulted in following groups: Transiently active (“0-1-0”), transiently suppressing (“1-0-1”), late active (“0-0-1”), late suppressing (“1-1-0”), prolonged active (“0-1-1”) and prolonged suppressing (“1-0-0”) reQTLs. The average of absolute eQTL-*β* and distribution of reQTL among these groups are shown in Fig. 2d (left panel). Of note, 83 reQTLs that were significant at all three time points (“1-1-1”) but with significant changes of the eQTL effect size are not illustrated and were excluded from the following analysis.

To further examine if eQTL-*β* and differential expression (DE) of the eQTL gene are congruent, DE between baseline and 90 min stimulation (∆_90min-baseline_) and DE between 90 min and 6 h stimulation (∆_6h-90min_) were calculated using limma and significant Δ_90min-baseline_ (*p* < 0.01) was encoded in binary (0;1) whereas significant Δ_6h-90min_ was encoded as “not significant” (0), “significant” (1), “significant, but opposite direction of Δ_90min-baseline_” (2). To determine the proportion of reQTL genes with congruent DE we quantified for transiently active/suppressing reQTLs the proportion of reQTL genes with significant Δ_90min-baseline_ and significant Δ_6h-90min_ with opposite direction (“1-2”), for late active/suppressing reQTLs we quantified the proportion of reQTL genes with not significant Δ_90min-baseline_ and significant Δ_6h-90min_ (“0-1”) and for prolonged active/suppressing reQTLs we quantified the proportion of reQTL genes with significant Δ_90min-baseline_ and either not significant Δ_6h-90min_ (expression stays the same) or significant Δ_6h-90min_ with same direction (fold change increases, “1-0” or “1-1”). To test if the proportion of reQTL genes with congruent DE was significantly enriched in each group (e.g. 37 congruent out of 71 transiently active reQTLs) we quantified the proportions of the same DE code (e.g. “1-2”) in the remaining groups (late active/suppressing and prolonged active/suppressing) and tested the proportions using Fisher’s exact test.

### Enrichment of functional annotations and fine mapping

We used the fgwas^7^ software to investigate the extent to which reQTLs and ceQTLs were enriched within specific annotation categories. Annotation information used by fgwas was derived from CADD variant consequence annotation^34^ (14 annotations) and Monocyte-specific annotations from Ensembl Regulatory build^35^ (6 annotations). To identify the set of annotations that would best fit the model, we first tested each of the 20 annotations in a joint data set of reQTLs and ceQTLs including distance to TSS in the analysis. 16 annotations individually improved the model likelihood but as many of these annotations are correlated with one another we used a stepwise selection approach to identify a final best-fitting model that included 13 annotations asterisked in Fig. 3b. We then ran fgwas including these 13 annotations for reQTLs and ceQTLs separately to estimate enrichment parameters and output reweighted summary statistics.

For each locus that contained at least one SNP with a posterior probability of association (PPA) > 0.3, we considered the SNP with the highest PPA from fgwas and tested the overlap of functional annotation sites of reQTL vs. ceQTLs using Fisher’s exact test. To increase power of reQTLs/ceQTLs overlapping functional annotation sites we mapped eQTLs using the mean of gene expression across all seven conditions. Fgwas steps were repeated as described above. Estimated enrichment parameters showed similar results and indicate the robustness of our analysis (Supplementary Fig. 8b).

### Natural selection analysis

We used two metrics, iHS and SDS, which detect signals of positive selection. The integrated haplotype score (iHS) measures the degree of extended haplotype homozygosity of the putatively selected allele over that of the putatively neutral allele^10^. iHS were calculated with the program selscan v1.1.0b^36^ with default parameters. We defined high iHS values as |iHS| > 1.5 in the CEU population. Furthermore, we used the recently published singleton density score (SDS)^11^, which detects very recent changes in allele frequencies from contemporary genome sequences. Publicly available SNP level SDS scores calculated from the UK10K Project reflect allele frequency changes during the past ~2000 to 3000 years in modern Britons, who are closely related to the German population^37^. We therefore applied these SDS scores to our cohort.

For each statistic (iHS, SDS), we determined the strongest signal of selection of all SNPs in high LD (*r^2^* > 0.8) with the best eQTL/ceQTL/reQTL SNP per gene. To assess significance, we then compared for each eQTL set the proportion of SNPs with |iHS| > 1.5 with the expected distribution obtained from re-sampling 10,000 sets of random SNPs matched for MAF and the number of SNPs in LD using the same parameters as described in Quach et al.^6^ using bins of MAF of 0.05 and LD bins of 0-2, 3-5, 6-10, 11-20, 21-50, and > 50 SNPs with *r^2^* > 0.8). Similarly, for SDS, we compared the median of SDS scores of eQTLs/ceQTLs/reQTLs, to the expected distribution obtained from resampling 10,000 sets of random SNPs matched for MAF and LD patterns.

To determine the effect of the derived allele on the immune response, we tested the proportion of reQTLs where the derived allele causes an increase versus decrease in response amplitude compared to the ancestral allele (Fig. 3d). reQTLs with increased activity include both reQTLs where the derived allele amplifies the induction of a gene or amplifies the suppression of a gene, whereas reQTLs with decreased activity will either reduce the induction of a gene or reduce the suppression of a gene. Overrepresentation of reQTLs with increased activity was evaluated using a binomial test.

### Colocalization analysis

Colocalization analysis was conducted using the R package coloc^12^. The method requires summary statistics for each SNP, which were summarized in Pickrell et al.^38^ or downloaded from ImmunoBase (http://www.immunobase.org) along with our eQTL data. A list of GWAS traits used in this analysis is provided in Supplementary Table 3. Coloc uses summary data from eQTL and GWAS studies in a Bayesian framework to identify GWAS signals that colocalize with eQTLs. We ran coloc using default parameter settings and a colocalization prior *p_12_* = 10^−6^. Coloc estimates posterior probability of association for either trait (PP0), association with gene expression (PP1), association with the trait (PP2), association with both phenotypes but distinct causal variants (PP3) and association with both phenotypes sharing the same causal variant (PP4). Regions with evidence for colocalization between gene expression and trait were defined as PP3 + PP4 ≥ 0.90 and PP4/PP3 ≥ 3 similar to what has been proposed by Guo et al.^39^ and are illustrated in Fig. 4a.

As eQTL summary statistics in the coloc analysis, we used two approaches to maximize our discovery power. First, from each locus we used the summary statistics of the condition with the strongest p-value. This is expected to provide robust discovery even in highly condition-specific loci. Furthermore, we also ran coloc with eQTLs mapped using the mean of gene expression across all seven conditions, which is expected to improve power when the eQTL signal is present in many conditions. All coloc results with PP3 + PP4 ≥ 0.90 are reported in Supplementary Table 3.

### Overlap between reQTLs and GWAS catalog

To assess the overlap between reQTLs and trait-associated variants, we downloaded the NHGRI-EBI GWAS Catalog (version 1.0.1, downloaded 2016/06/14). A reported GWAS SNP was considered to coincide with an reQTL if the GWAS SNP was in high LD (*r^2^* >0.8) with the lead eSNP per gene. A full list of these GWAS reQTLs is provided in Supplementary Table 4.

### Estimating the relative contribution of reQTLs and ceQTLs on immune-mediated traits

We used the fgwas^7^ software to investigate the extent to which reQTLs and ceQTLs were enriched in risk loci of immune-mediated traits, following the approach of Li et al.. A list of GWAS traits used in this analysis is provided in Supplementary Table 3. Due to the limited number of 417 reQTLs and 677 ceQTLs, we loosened the eQTL cutoffs for reQTLs and ceQTLs. For reQTLs, we considered all reQTLs that were significant after Benjamini-Hochberg FDR 5% correction (instead of Bonferroni correction), which resulted in 1128 reQTLs. For ceQTLs, we considered all ceQTLs with *p*_beta_ > 0.005 when testing each of the six stimulated conditions with the baseline condition, which resulted in 1165 ceQTLs. For both eQTLs, all associations with *p* < 10^−4^ were used as input, and fgwas analysis was performed for reQTLs and ceQTLs separately. Of note, this analysis was robust to different eQTL association p-value cutoffs (*p* < 10^−4^, 10^−5^, 10^−6^) suggesting that the enrichment is not simply due to the power of detection (**Supplementary Fig. 10, Supplementary Fig. 11**).

### Data availability

Full summary statistics of the eQTL analysis, gene expression and genotyping data will be available at ArrayExpress and the European Genome-phenome Archive [to be released at publication]. In addition to results tables for all seven conditions provided in Supplementary Table 2, all eQTL results are available in the ImmunPop QTL browser (http://immunpop.com/kim/eQTL), which provides multiple interactive visualization and data exploration features for eQTLs.

## Acknowledgements

We thank all blood volunteers for participating to this study. We acknowledge our laboratory technicians and colleagues responsible for database management. S.K.-H. is supported by a research fellowship of the DFG. J.S. and V.H. received support for this work from the BONFOR research program, individual grant O-149.0094. JS was supported by the NIH/DFG Research Career Transition Award. M.M.N. received support for this work from the Alfried Krupp von Bohlen und Halbach-Stiftung. V.H. is supported by the European Research Council (ERC 243046). M.M.N. and V.H. are members of the DFG funded Excellence Cluster ImmunoSensation. P.B. was supported by the SFB/Transregio 57 (TP25 and Q1) and an individual DFG grant (BO 3755/1-1). T.L. and P.M. were supported by the NIH grant R01MH106842. T.L. was supported by the NIH grants UM1HG008901 and 1U24DK112331-01. T.L. and S.E.C were supported by the NIH contract HHSN2682010000029C.

## Author Contributions

S.K.-H., J.S., V.H. initiated the study. S.K.-H., B.P., P.M., Y.N., N.G., S.C., L.B.B., J.K.P., B.M.-M., J.S., V.H., T.L. analyzed and interpreted the data. S.K.-H., J.B., M.B., V.K., E.B., N.F., P.B. performed the molecular genetic experiments. S.K.-H., M.B. characterized the volunteers and collected blood samples. S.K.-H., M.M.N., J.S., V.H., T.L., prepared the manuscript, with feedback from the other authors.

## References

1. Kim, S. et al. Characterizing the genetic basis of innate immune response in TLR4-activated human monocytes. Nat Commun 5, 5236 (2014).

2. Barreiro, L. B. et al. Deciphering the genetic architecture of variation in the immune response to Mycobacterium tuberculosis infection. Proc. Natl. Acad. Sci. U.S.A. 109, 1204–1209 (2012).

3. Lee, M. N. et al. Common Genetic Variants Modulate Pathogen-Sensing Responses in Human Dendritic Cells. Science 343, 1246980–1246980 (2014).

4. Fairfax, B. P. et al. Innate Immune Activity Conditions the Effect of Regulatory Variants upon Monocyte Gene Expression. Science 343, 1246949–1246949 (2014).

5. Nédélec, Y. et al. Genetic Ancestry and Natural Selection Drive Population Differences in Immune Responses to Pathogens. Cell 167, 657–664.e21 (2016).

6. Quach, H. et al. Genetic Adaptation and Neandertal Admixture Shaped the Immune System of Human Populations. Cell 167, 643–649.e17 (2016).

7. Pickrell, J. K. Joint Analysis of Functional Genomic Data and Genome-wide Association Studies of 18 Human Traits. The American Journal of Human Genetics 94, 559–573 (2014).

8. Ostuni, R. et al. Latent Enhancers Activated by Stimulation in Differentiated Cells. Cell 152, 157–171 (2013).

9. Aguet, F. et al. Local genetic effects on gene expression across 44 human tissues. bioRxiv 1–24 (2016). doi:10.1101/074450

10. Voight, B. F., Kudaravalli, S., Wen, X. & Pritchard, J. K. A map of recent positive selection in the human genome. PLoS Biol. 4, e72 (2006).

11. Field, Y. et al. Detection of human adaptation during the past 2000 years. Science 354, 760–764 (2016).

12. Giambartolomei, C. et al. Bayesian test for colocalisation between pairs of genetic association studies using summary statistics. PLoS Genet 10, e1004383 (2014).

13. Teslovich, T. M. et al. Biological, clinical and population relevance of 95 loci for blood lipids. Nature 466, 707–713 (2010).

14. Dubois, P. C. A. et al. Multiple common variants for celiac disease influencing immune gene expression. Nat. Genet. 42, 295–302 (2010).

15. Schizophrenia Working Group of the Psychiatric Genomics Consortium. Biological insights from 108 schizophrenia-associated genetic loci. Nature 511, 421–427 (2014).

16. Perry, J. R. B. et al. Parent-of-origin-specific allelic associations among 106 genomic loci for age at menarche. Nature 514, 92–97 (2014).

17. Welter, D. et al. The NHGRI GWAS Catalog, a curated resource of SNP-trait associations. Nucleic Acids Res 42, D1001–6 (2014).

18. Li, Y. I. et al. RNA splicing is a primary link between genetic variation and disease. Science 352, 600–604 (2016).

19. Bentham, J. et al. Genetic association analyses implicate aberrant regulation of innate and adaptive immunity genes in the pathogenesis of systemic lupus erythematosus. Nat. Genet. 47, 1457–1464 (2015).

20. Lappalainen, T. et al. Transcriptome and genome sequencing uncovers functional variation in humans. Nature 501, 506–511 (2013).

21. Alasoo, K. et al. Genetic effects on chromatin accessibility foreshadow gene expression changes in macrophage immune response. bioRxiv 1–39 (2017). doi:10.1101/102392

22. Li, X. et al. The impact of rare variation on gene expression across tissues. bioRxiv 1–22 (2016). doi:10.1101/074443

23. Astle, W. J. et al. The Allelic Landscape of Human Blood Cell Trait Variation and Links to Common Complex Disease. Cell 167, 1415–1429.e19 (2016).

24. Johnson, W. E., Li, C. & Rabinovic, A. Adjusting batch effects in microarray expression data using empirical Bayes methods. Biostatistics 8, 118–127 (2006).

25. Leek, J. T., Johnson, W. E., Parker, H. S., Jaffe, A. E. & Storey, J. D. The sva package for removing batch effects and other unwanted variation in high-throughput experiments. Bioinformatics 28, 882–883 (2012).

26. Smyth, G. K. Linear Models and Empirical Bayes Methods for Assessing Differential Expression in Microarray Experiments. Statistical Applications in Genetics and Molecular Biology 3, 1–25 (2011).

27. Falcon, S. & Gentleman, R. Using GOstats to test gene lists for GO term association. Bioinformatics 23, 257–258 (2007).

28. O’Connell, J. et al. A General Approach for Haplotype Phasing across the Full Spectrum of Relatedness. PLoS Genet 10, e1004234–21 (2014).

29. Howie, B. N., Donnelly, P. & Marchini, J. A flexible and accurate genotype imputation method for the next generation of genome-wide association studies. PLoS Genet 5, e1000529 (2009).

30. Consortium, T. 1. G. P. et al. An integrated map of genetic variation from 1,092 human genomes. Nature 490, 56–65 (2012).

31. Ongen, H., Buil, A., Brown, A. A., Dermitzakis, E. T. & Delaneau, O. Fast and efficient QTL mapper for thousands of molecular phenotypes. Bioinformatics 32, 1479–1485 (2016).

32. Szklarczyk, D. et al. STRING v10: protein-protein interaction networks, integrated over the tree of life. Nucleic Acids Res 43, D447–D452 (2015).

33. Storey, J. D. & Tibshirani, R. Statistical significance for genomewide studies. Proc Natl Acad Sci USA 100, 9440–9445 (2003).

34. Kircher, M. et al. A general framework for estimating the relative pathogenicity of human genetic variants. Nat. Genet. 46, 310–315 (2014).

35. Zerbino, D. R., Wilder, S. P., Johnson, N., Juettemann, T. & Flicek, P. R. The ensembl regulatory build. Genome Biol. 16, 56 (2015).

36. Szpiech, Z. A. & Hernandez, R. D. selscan: an efficient multithreaded program to perform EHH-based scans for positive selection. Molecular Biology and Evolution 31, 2824–2827 (2014).

37. Lappalainen, T. et al. Genomic landscape of positive natural selection in Northern European populations. European Journal of Human Genetics 18, 471–478 (2009).

38. Pickrell, J. K. et al. Detection and interpretation of shared genetic influences on 42 human traits. Nat. Genet. 48, 709–717 (2016).

39. Guo, H. et al. Integration of disease association and eQTL data using a Bayesian colocalisation approach highlights six candidate causal genes in immune-mediated diseases. Human Molecular Genetics 24, 3305–3313 (2015).

